# Germ cells do not progress through spermatogenesis in the infertile zebrafish testis

**DOI:** 10.1101/2023.09.05.556432

**Authors:** Andrea L. Sposato, Darren R. Llewellyn, Jenna M. Weber, Hailey L. Hollins, Madison N. Schrock, Jeffrey A. Farrell, James A. Gagnon

## Abstract

Vertebrate spermatogonial stem cells maintain sperm production over the lifetime of an animal but fertility declines with age. While morphological studies have greatly informed our understanding of typical spermatogenesis, the molecular and cellular mechanisms underlying spermatogenesis are not yet understood, particularly with respect to the onset of fertility. We used single-cell RNA sequencing to generate a developmental atlas of the zebrafish testis. Using 5 timepoints across the adult life of a zebrafish, we described cellular profiles in the testis during and after fertility. While all germ cell stages of spermatogenesis are detected in testes from fertile adult zebrafish, testes from older infertile males only contained spermatogonia and a reduced population of spermatocytes. These remaining germ cells are transcriptionally distinct from fertile spermatogonia. Immune cells including macrophages and lymphocytes drastically increase in abundance in infertile testes. Our developmental atlas reveals the cellular changes as the testis ages and defines a molecular roadmap for the regulation of male fertility.

## Introduction

The vertebrate testis produces sperm throughout the fertile lifetime of the animal from a population of spermatogonial stem cells. During spermatogenesis, differentiating germ cells undergo transcriptional reprogramming as they mature through a diversity of cell types (Soumillon et al. 2013, Melé et al. 2015). Somatic cells of the testis support spermatogenesis by providing a conducive environment for a balance of germ cell proliferation and differentiation. Several imaging-based strategies have captured the morphology of developing germ and somatic cells, providing insight into the differences in testis composition and organization across species (Maack and Segner 2003, Schulz et al. 2005, Wang et al. 2007, Leal et al. 2009, Uribe et al. 2014, Lee et al. 2017, de Siqueira-Silva et al. 2019).

Mammals and zebrafish share an overall testis architecture composed of several tightly coiled seminiferous tubules (Schulz et al. 2010). As in most animals, spermatogenesis is maintained throughout most of zebrafish adulthood. Unlike mammals, germ cells in zebrafish are not in direct contact with the basement membrane surrounding tubules. Instead, spermatogenesis is a cystic process whereby Sertoli cells surround individual undifferentiated spermatogonia and support their differentiation throughout spermatogenesis (Schulz et al. 2005). Within a Sertoli cyst, germ cells first develop in a clonal syncytium of type A spermatogonia before undergoing 9 mitotic divisions as type B spermatogonia then entering meiosis (Leal et al. 2009). After meiosis, germ cells enter the spermiogenic phase, where spermatids develop into spermatozoa following nuclear condensation, organelle elimination, and formation of the flagellum. The cyst then opens to release mature spermatozoa in the lumen of seminiferous tubules (Schulz et al. 2010). In addition to Sertoli cells, other somatic cell types also serve important roles during spermatogenesis. Leydig cells produce testosterone and insulin-like 3 which promote spermatogenesis via Sertoli cells (Walker 2010, Crespo et al. 2021). A host of immune cells are also present in the testis, including macrophages and various lymphocytes. Regulatory T cells play an important role in immune homeostasis during zebrafish testis development (Li et al. 2020). In mammalian testes, macrophage and lymphocyte populations have also been characterized (Guo et al. 2020, Bhushan et al. 2020) and inflammation has been described as a signature of testicular aging (Matzkin et al. 2016, Nie et al. 2022). To date, the full scope of cell type diversity and plasticity during zebrafish spermatogenesis and aging is not well understood.

Zebrafish become fertile at 3 months old and maintain gametogenesis until about 2 years of age when they are unable to breed or make sperm. We do not know how the cell type composition and transcriptional profiles of the testis may change throughout fertility and the transition to infertility. Single-cell RNA sequencing (scRNAseq) has been used to describe the molecular and cellular composition of testes from several vertebrate and invertebrate organisms. A recent scRNAseq survey profiled the cell types present in 5 month old zebrafish (Qian et al. 2022). As expected, these testes exhibited a continuum of cell types from spermatogonia to elongated spermatids. However, this stage of life marks just the sunrise of a dynamic and continually maintained process throughout the adult life of a zebrafish. Another study compared 18-month testes from control zebrafish and infertile fish that were exposed to an endocrine-disruptive chemical early in life (Haimbaugh et al. 2022). Notably, endocrine disruption caused arrest or apoptosis of post-meiotic germ cells after typical development through spermatogonial and meiotic phases. These results suggest that meiosis acts as a key checkpoint for sperm development. However, profiles of somatic cells are missing from this atlas, and it remains unknown whether this drug-induced post-meiotic apoptosis/arrest is also present in natural cases of infertility. While single timepoint atlases of the zebrafish testis provide snapshots of testicular cell type composition, to probe the mechanisms that control infertility we need to capture the dynamic changes in the testis as fertility declines.

In this study, we describe a developmental atlas of the whole zebrafish testis which characterizes cell types present throughout the fertility window and beyond. We used scRNAseq to profile cell types, enabling us to identify all stages of germ cell differentiation and somatic cells of the testis including Sertoli, Leydig, vascular smooth muscle cells, and immune cells such as macrophages, T cells, and natural killer cells. In naturally infertile testes, we identify a larger and more diverse population of immune cells and an absence of post-meiotic germ cells. While infertile testes are not depleted of germ cells, these germ cells adopt a distinct transcriptional state and do not develop beyond meiotic stages. Our developmental atlas provides insights regarding the maintenance and breakdown of fertility with cellular and molecular resolution.

## Results

### Timecourse scRNAseq identifies cell types present in the zebrafish testis throughout fertility

To define the cellular composition of zebrafish testes during fertility, we dissected whole testes from fertile adult zebrafish at four ages (5, 12, 20, and 22 months old), dissociated whole tissue, and profiled each sample by scRNAseq (Fig. 1A, S1A). Representative histological images of fertile zebrafish testes show densely packed seminiferous tubules with germ cells proceeding through spermatogenesis and somatic support cells in the interstitial spaces (Fig. 1B, S2). Rates of transcription are highest in the testis compared to every other organ (Xia et al. 2020), so we filtered for high quality transcriptomes based on the percentage of mitochondrial reads (<5%) and the number of genes detected per cell using a threshold established for each data set based on the interquartile range (Fig. S1C). Next, we integrated the six biological samples from all four timepoints to establish a composite atlas of the fertile zebrafish testes that contains 32,659 cells (Fig. 1C, S1B).

**Figure 1.**
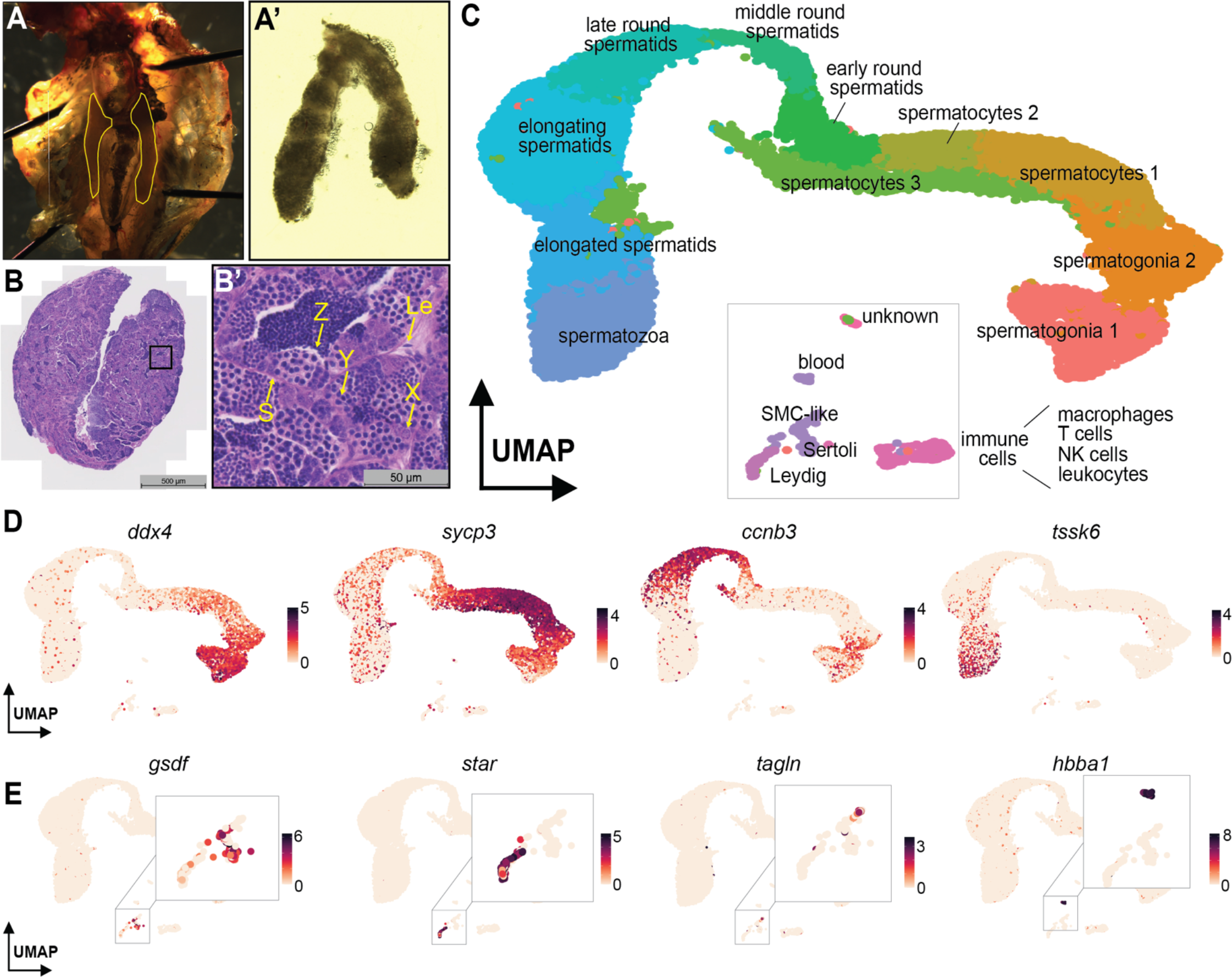
Single-cell RNA sequencing of 32,659 cells from six fertile adult zebrafish testes. **(A-A’)** Testes dissected from fertile 12 month old zebrafish. Testes are highlighted in yellow in the trunk image (A). **(B-B’)** Histology of adult testes from a 12 month old zebrafish. Yellow arrows point to basement membrane of seminiferous tubule (S), undifferentiated spermatogonia (X), spermatocytes (Y), spermatids (Z), and Leydig cell (Le). **(C)** An atlas of fertile zebrafish testes. Cell clusters were identified using canonical markers. **(D)** Representative markers of spermatogonia, spermatocytes, spermatids and spermatozoa. **(E)** Representative markers of Sertoli, Leydig, smooth muscle cells, and blood.

We identified all cell types within this fertile atlas using marker genes (Table 1, Fig S1D). We identified germ cells as they progressed through all stages of differentiation from early spermatogonia to spermatozoa (Fig. 1D). We also discovered small populations of somatic cells such as Sertoli, Leydig, smooth muscle-like cells and various immune cells (Fig. 1C, E). Overall, >90% of cells in the atlas were differentiating germ cells, which is likely an overrepresentation of their relative abundances in intact tissue. We hypothesize that the enrichment of germ cells may be attributed to dissociation sensitivity, where irregular shape and large size of somatic cell types may have decreased their capture rate during 10X scRNAseq. Notably, every cluster is composed of cells from all four sample ages with the exception of cluster 14 which is unique to the 5 month atlas (Fig. S1E). Top differentially expressed genes in this cluster are associated with cell types found in the liver, suggesting this cluster is derived from contamination during dissection. We conclude that our composite atlas captures all known testis cell types and can serve as a cell type reference for this tissue. A web-based application for exploring the single-cell datasets presented in this paper is available at https://github.com/asposato/zebrafish_testis_fertility.

**Table 1.** A subset of marker genes used to identify testicular cell types.

| cell type | marker | reference | DOI |
| --- | --- | --- | --- |
| germ cells | *ddx4* | Yoon et al., 1997 | 10.1242/dev.124.16.3157 |
|  | *piwil1* | Houwing et al., 2007 | 10.1016/j.cell.2007.03.026 |
|  | *dmrt1* | Webster et al., 2017 | 10.1016/j.ydbio.2016.12.008 |
| spermatogonia | *sumo3b* | Qian et al., 2022 | 10.3389/fgene.2022.851719 |
|  | *pcna* | Ye et al., 2023 | 10.3389/fendo.2023.1044318 |
|  | *hist1h4l* | Qian et al., 2022 | 10.3389/fgene.2022.851719 |
|  | *ccnb3* | Ozaki et al., 2011 | 10.1016/j.gep.2011.03.002 |
| spermatocytes | *sycp3* | Ozaki et al., 2011 | 10.1016/j.gep.2011.03.002 |
|  | *dmc1* | Beer & Draper, 2012 | 10.1016/j.ydbio.2012.12.003 |
|  | *spo11* | Zhang et al., 2020 | 10.1111/are.14829 |
|  | *e2f5* | Xie et al., 2020 | 10.1371/journal.pgen.1008655 |
| spermatids | *ccnb3* | Ozaki et al., 2011 | 10.1016/j.gep.2011.03.002 |
|  | *larp1b* | Qian et al., 2022 | 10.3389/fgene.2022.851719 |
|  | *edrf1* | Qian et al., 2022 | 10.3389/fgene.2022.851719 |
| spermatozoa | *tssk6* | Li. et al., 2011 | 10.1093/molehr/gaq071 |
|  | *spag8* | Wu et al., 2010 | 10.1016/j.febslet.2010.05.016 |
| Sertoli | *gsdf* | Gautier et al., 2011 | 10.1095/biolreprod.111.091892 |
|  | *krt18a.1* | Qian et al., 2022 | 10.3389/fgene.2022.851719 |
| Leydig | *star* | Gautier et al., 2011 | 10.1095/biolreprod.111.091892 |
|  | *cyp17a2* | Gautier et al., 2011 | 10.1095/biolreprod.111.091892 |
| SMC-like | *tagln* | Santoro et al., 2009 | 10.1016/j.mod.2009.06.1080 |
|  | *acta2* | Georgijevic et al., 2007 | 10.1002/dvdy.21165 |
|  | *dcn* | Järveläinen et al., 2015 | 10.1016/j.matbio.2015.01.023 |
| blood | *hbba1* | Qian et al., 2022 | 10.3389/fgene.2022.851719 |
|  | *fli1b* | Craig et al. 2015 | 10.1161/ATVBAHA.114.304768 |
| leukocytes | *coro1a* | Song et al., 2004 | 10.1073/pnas.0407241101 |
|  | *ptprc* | Rougeot et al., 2019 | 10.3389/fimmu.2019.00832 |
| macrophages | *mpeg1.1* | Kuil et al., 2020 | 10.7554/eLife.53403 |
|  | *csf1ra* | Kuil et al., 2020 | 10.7554/eLife.53403 |
| T cells | *zap70* | Yoon et al., 2015 | 10.1371/journal.pone.0126378 |
|  | *cd4-1* | Yoon et al., 2015 | 10.1371/journal.pone.0126378 |
|  | *runx3* | Kalev-Zylinska et al., 2003 | 10.1002/dvdy.10388 |
|  | *foxp3a* | Li. et al., 2020 | 10.1016/j.jgg.2020.07.006 |
| neutrophils | *mpx* | Mathias et al., 2006 | 10.1189/jlb.0506346 |
|  | *lyz* | Meijer et al., 2007 | 10.1016/j.dci.2007.04.003 |
| B cells | *pax5* | Roessler et al., 2007 | 10.1128/MCB.01192-06 |
| NK cells | *nkl.2* | Pereiro et al., 2015 | 10.1016/j.dci.2015.03.009 |

### Testes from infertile zebrafish lack fully differentiated germ cells

A scRNAseq atlas of the infertile zebrafish testis would permit comparisons to uncover the mechanisms that regulate fertility. In laboratory settings, domesticated zebrafish can live for 3-5 years but fecundity decreases around 2 years of age. Testes of infertile animals are smaller and disorganized with expanded interstitial spaces and germ cells in less dense clusters (Fig. 2A, B). Representative images of infertile zebrafish demonstrate that the remaining germ cells within seminiferous tubules are mostly at earlier stages of spermatogenesis and interstitial spaces are expanded between tubules (Fig. S3). We dissected testes from two 27 month old zebrafish, dissociated whole tissue, and profiled each sample by scRNAseq. After filtering, the resulting infertile zebrafish testis atlas contained 24,469 cells (Fig. 2C, S4A).

**Figure 2.**
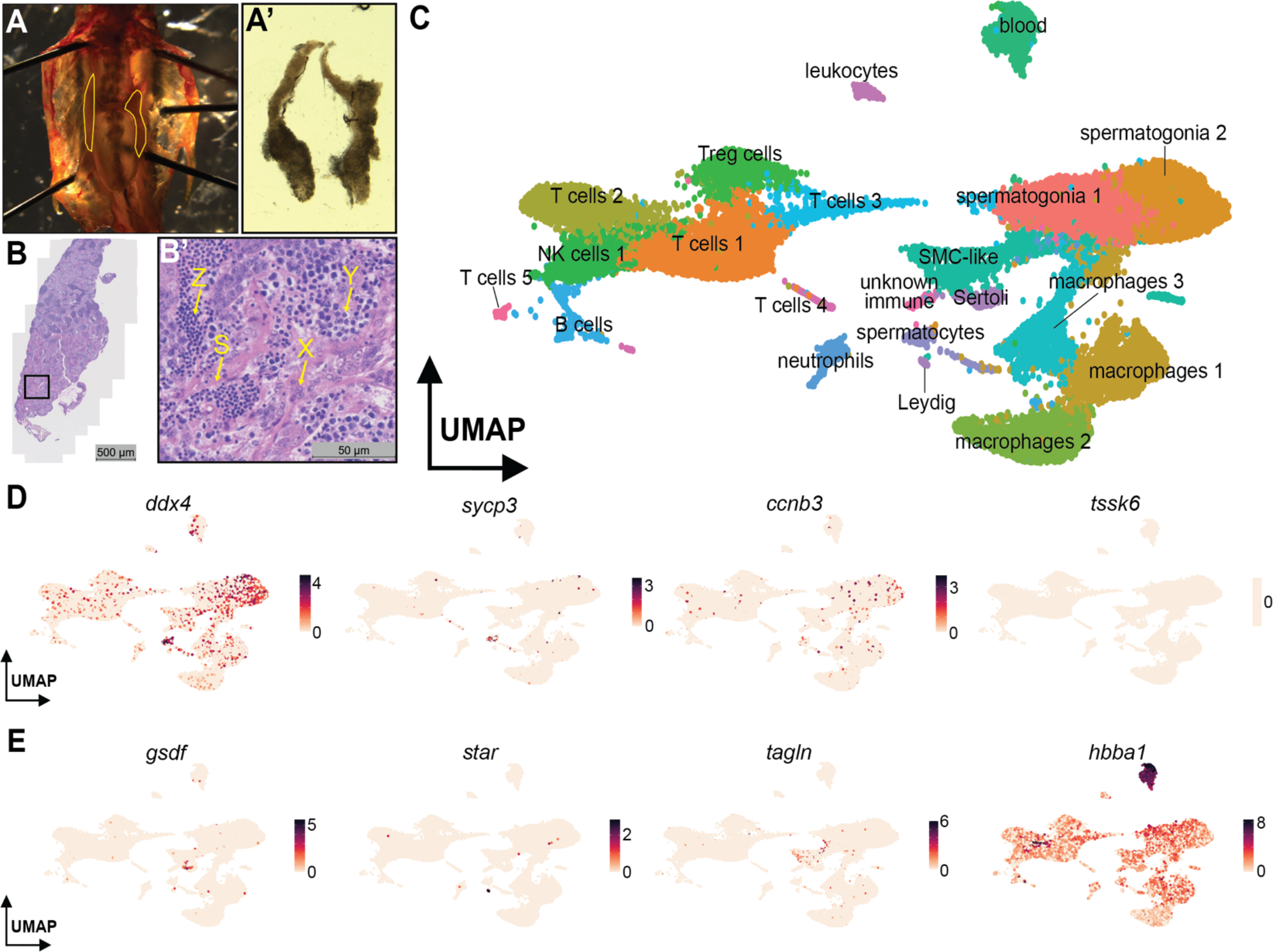
Single-cell RNA sequencing of 24,469 cells from two infertile 27 month old adult zebrafish testes. **(A-A’)** Testes dissected from infertile 26 month old zebrafish. Testes are highlighted in yellow in the trunk image. **(B-B’)** Histology of adult testes from a 26 month old zebrafish. Yellow arrows point to basement membrane of seminiferous tubule (S), undifferentiated spermatogonia (X), spermatocytes (Y) and spermatids (Z). **(C)** An atlas of infertile zebrafish testes. Cell clusters were identified using canonical markers. **(D)** Representative markers of spermatogonia and spermatocytes. Cyclin B3 (*ccnb3*) is expressed by round spermatids and 16-cell stage spermatogonial cysts. **(E)** Representative markers of Sertoli, Leydig, smooth muscle cells, and blood.

We again identified all cell types within this atlas using marker genes (Fig. 2C, D, S4B). In the infertile atlas, only three clusters of germ cells were identified while the remaining 17 clusters were somatic cell types. While 93% of the cells in the composite fertile atlas are germ cells at various stages of development, germ cells make up only 32% of the infertile atlas. Of the germ cell clusters, marker genes for spermatogonia and spermatocytes are detected, yet post-meiotic germ cells were absent (Fig. 2D). Instead, somatic cell types make up the majority of the infertile atlas with immune cells accounting for much of this increase.

### Somatic cell type composition shifts in the infertile testis

We compared somatic cell types between fertile and infertile testes to discover changes that may correlate with infertility. Coordination among immune cells is critical to the establishment and maintenance of tissue homeostasis (Bhushan et al. 2020, Fan et al. 2021). To investigate changes in immune cell composition, we subsetted immune cells from fertile and infertile testes using marker genes and reclustered these cells to create an immune atlas of the zebrafish testis (Fig. 3A, S5A). We identified cell types within this immune atlas using marker genes (Table 1, Fig. 3B, S5B). We found six T cell subtypes, natural killer cells, B cells, four macrophage subtypes, neutrophils, and one unknown leukocyte cell type. Most cells are from the infertile sample, although there was representation of all cell types within the fertile testes (Fig 3C, 5SA). We categorized cell types into broad somatic categories and calculated the relative abundance for each age. Many somatic cell types are elevated in infertile testes, particularly in macrophage and lymphocyte populations (Fig. 3D).

**Figure 3.**
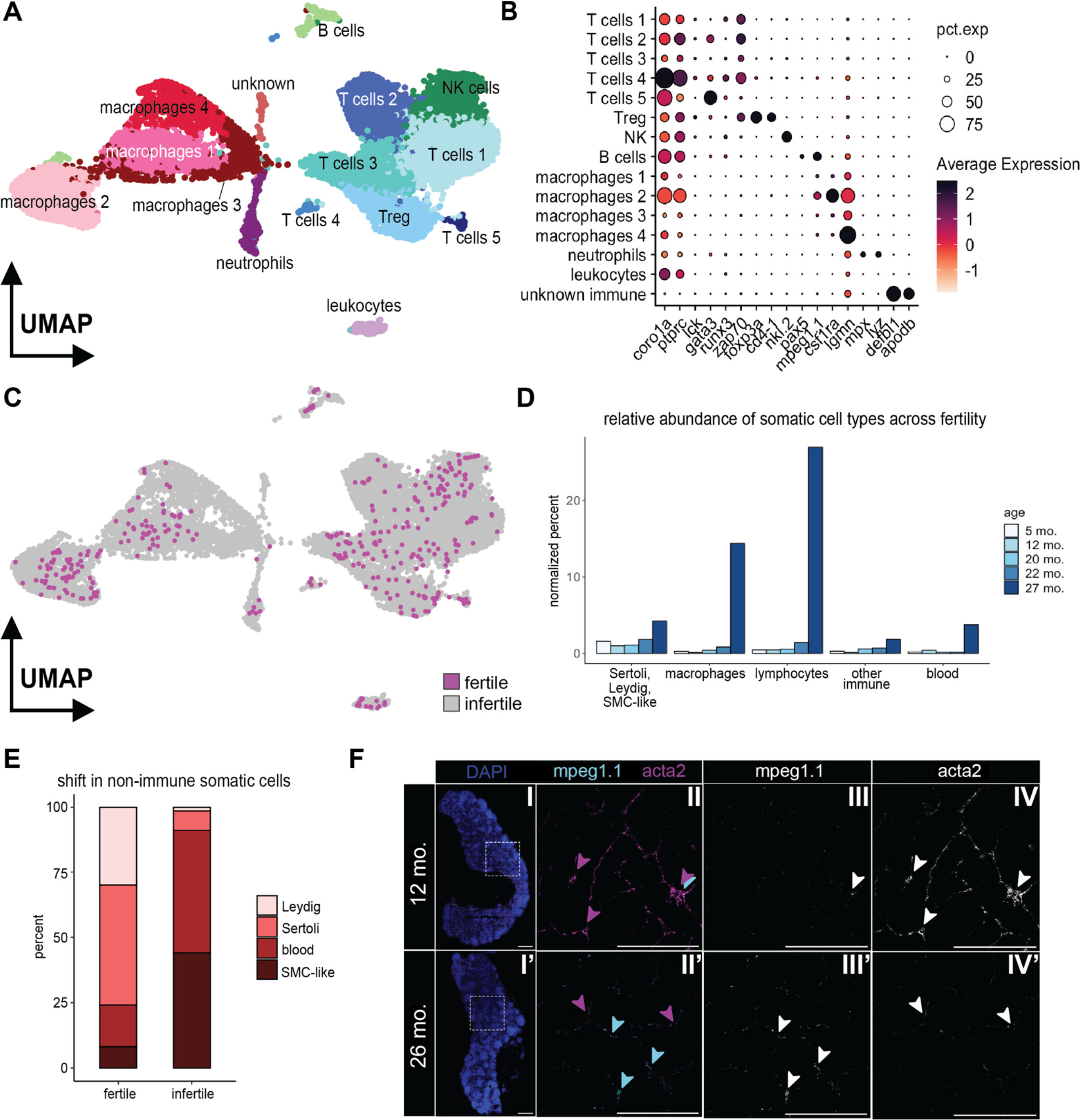
Reclustered immune cells from fertile and infertile atlases. **(A)** The infertile testes are enriched for immune cells. Particularly, large populations of T cells and macrophages were detected. **(B)** Canonical marker genes were used to identify each cell type. **(C)** Cells from fertile animals highlighted on immune cells atlas. **(D)** Relative abundance of somatic cell types from each sample age. **(E)** Proportions of Sertoli, Leydig, blood and smooth muscle in fertile and infertile testes. **(F)** RNAscope of smooth muscle marker *acta2* and macrophage marker *mpeg1*.*1* in fertile and infertile testes. (I-I’) DAPI staining of testis lobes from 12 mo. and 26 mo. zebrafish. Square inset used for images in panels II-IV. (I-II’) Merge of *acta2* (magenta) and *mpeg1*.*1* (cyan) markers. (III-III’) Macrophage marker *mpeg1*.*1*. (IV-IV’) Smooth muscle marker *acta2*. Arrowheads highlight examples of fluorescence.

We also observed an expansion of some non-immune somatic cell populations in infertile testes (Fig. 3D). Split by cell type, blood cells and SMC-like cells were 9-fold higher in abundance in infertile testes, while Leydig and Sertoli cells proportionally decreased in abundance (Fig. 1E). Histological images (Fig. 1A-B, S2, 2A-B, S3) also show a large expansion of the interstitial spaces around tubules in infertile testes. To confirm changes in somatic cell abundances in the zebrafish testis, we used RNA in situ hybridization to detect macrophages and smooth muscle cells in both fertile and infertile testes. We found increased signal for the macrophage marker *mpeg1*.*1* and decreased signal for the smooth muscle marker *acta2* in infertile testes compared to fertile (Fig. 3E-F, S6A). Together, these data suggest that where germ cells are not progressing completely through spermatogenesis, the tubules give way to larger interstitial areas of connective tissue that contain many macrophages. Interestingly, while the proportion of blood and SMC-like cells seems to increase in infertile testes according to the single-cell data, molecular imaging shows that smooth muscle marker *acta2* diminishes in infertile testes. *Acta2* only weakly identifies SMC-like cells in the single-cell atlases. However, we do observe more abundant expression of *decorin* (*dcn*), a gene known to be expressed by stressed vascular endothelial cells and fibroblasts, in the SMC-like cells of the infertile testes (Fig. S6B, Järveläinen et al. 2015). The SMC-like cells in these interstitial areas may be breaking down between seminiferous tubules in aging testes.

### Spermatogenesis is stable and continuous through fertility but incomplete in older testes

The absence of later stages of spermatogenesis in testes from infertile zebrafish led us to hypothesize that there may be a developmental block at meiosis. The difference in tissue integrity and lack of densely packed germ cells in the infertile histological images further supported this hypothesis (Fig. 1A-B, S2, 2A-B, S3). As animals age, we hypothesized that the proportion of germ cells may shift toward more spermatogonia compared to spermatozoa. 27 month-old testes completely lack spermatozoa but contain more spermatogonia than 5 month-old testes (Fig. 4A). Next, we sought to quantify how the progression of germ cell differentiation changed as the fertile testis ages using URD pseudotime analysis (Fig. 4B, Farrell et al. 2018). In this analysis, 0 represents the most spermatogonial state, and 1 represents the most differentiated spermatid state. We split the integrated object by age to visualize the proportion of germ cells at each stage of spermatogenesis (Fig. 4C). While all fertile stages were similar, we observed a modest increase in spermatogonial proportion and a reciprocal decrease in contribution to later stages of spermatogenesis in older fertile testes (20 and 22 months) relative to testes from younger males (5 and 12 months). As animals approach infertility, our data suggests that the spermatogonia population declines in its ability to give rise to post-meiotic cells.

**Figure 4.**
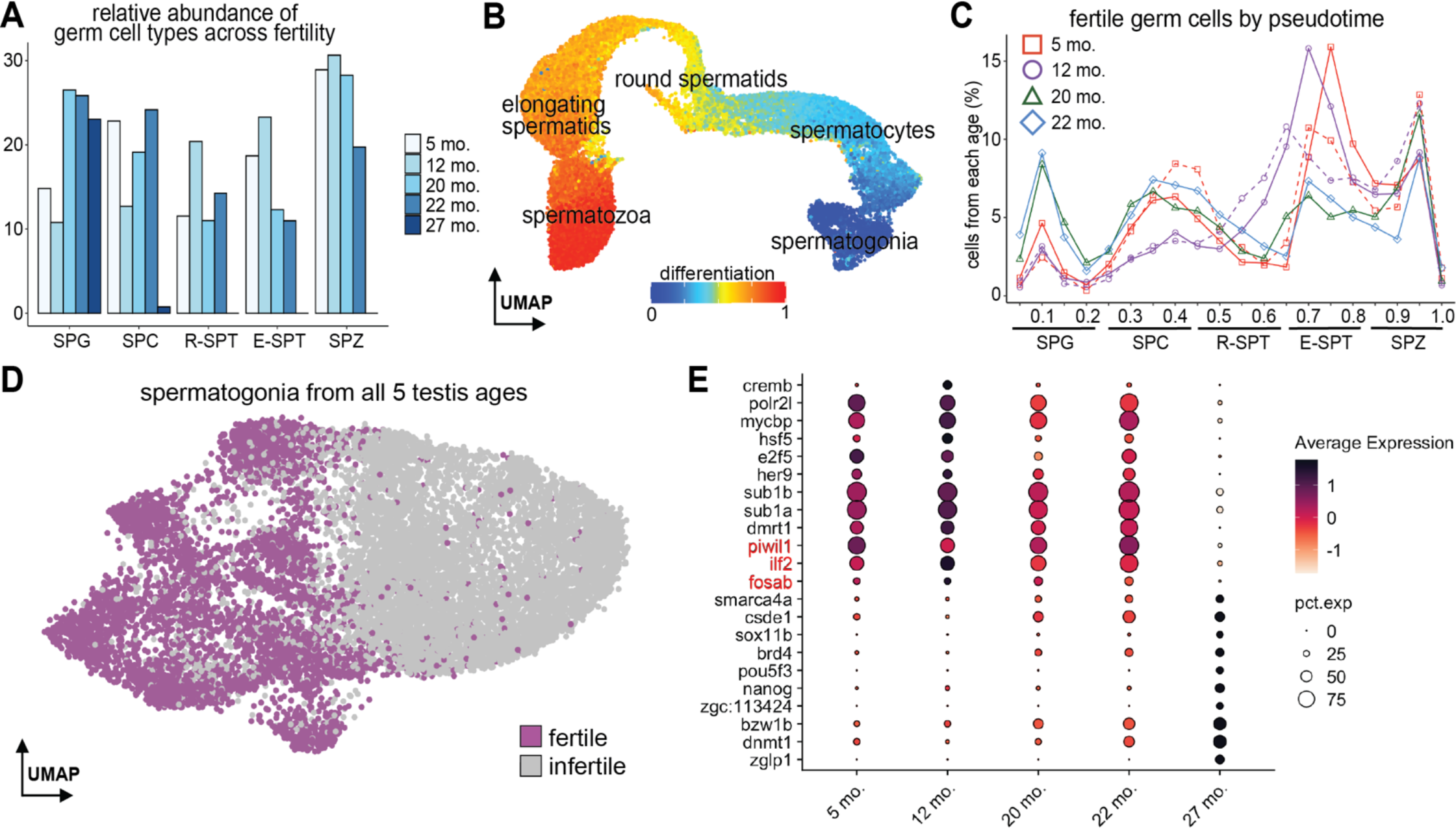
Spermatogenesis is stable and continuous through fertility. **(A)** Relative abundance of germ cell types from each sample age. **(B)** Fertile germ cells colored by pseudotime-determined scores of differentiation using URD. **(C)** Normalized percent of cells from each fertile age binned across differentiation. Ages with replicates are denoted with dashed lines. **(D)** Reclustered spermatogonia from fertile and infertile atlases. **(E)** Dotplot shows the top 10 transcription factors for fertile and infertile testes compared with known regulators (emphasized in red).

To investigate this phenomenon, we subsetted and reclustered spermatogonia from both fertile and infertile atlases. We found that the majority of germ cells from the infertile testes are transcriptionally distinct from fertile spermatogonia (Fig. 4D). Several known regulators of spermatogenesis such as *piwil1* and *ilf2* are more highly expressed in fertile germ cells compared to infertile (Fig. 4E). *E2f5*, recently identified as a new marker for undifferentiated spermatogonia (Xie et al. 2020, Qian et al. 2022), is highly expressed at 5 months, decreases in expression by 22 months, and is almost undetectable in spermatogonia from 27 month old testes. Perhaps the contrasting patterns of gene expression underlie the failure of spermatogonia in infertile testes to progress past meiosis. Together, our imaging and single-cell transcriptomic data suggest that infertile testes have a large population of spermatogonia and some spermatocytes but a dearth of post-meiotic cell types. We hypothesized that the expansion of immune cells we observe in infertile testes may help facilitate the apoptosis and removal of dying cells in the testis, perhaps explaining the absence of post-meiotic germ cells in infertile testes by a mechanism similar to what is observed in endocrine-disrupted testes (Haimbaugh et al. 2022). However, we did not detect differences in the expression of markers for apoptosis between fertile and infertile testes (Fig. S7), suggesting instead there may be a block at meiosis preventing the progression of germ cells past a critical checkpoint.

## Discussion

In this study, we used timecourse scRNAseq to profile cells of the aging zebrafish testis. Our approach captures the stability of spermatogenesis during fertility and the dynamic changes within the testis during the onset of infertility. Germ cells are relatively stable in fertile zebrafish of all ages with proportions of germ cells following a consistent pattern across differentiation. In older yet still fertile testes, we observed a shift in abundance toward less differentiated cell types, but cell types at later stages of spermatogenesis are still detected. However, in infertile testes we did not detect post-meiotic germ cells. Histology of fertile and infertile testes confirmed that seminiferous tubules of fertile testes are densely packed with germ cells across the spectrum of differentiation while tubules of infertile testes have noticeably fewer germ cells of the size and density of post-meiotic germ cells. Infertile testes also exhibit reduced structural integrity with expanded interstitial spaces throughout the tissue without maintenance or expansion of smooth muscle. Infertile testes contain spermatogonial stem cells, but they have a distinct transcriptional identity and spermatogenesis is interrupted during or following the meiotic phase. Our developmental atlases describe testis composition across the fertile and infertile lifespan of a zebrafish, providing a foundation for probing the mechanisms underlying infertility.

Our results also highlight a diversity of immune cells lacking from previous scRNAseq studies of zebrafish testes. While some immune cells are found in the fertile atlas, the immune cell population of the infertile atlas is more diverse and proportionally far larger than the germ cell population. Notably, lymphocyte and macrophage populations form the majority of cells in infertile testes. In a previous study of endocrine disrupted testes, spermatogenesis failure in post-meiotic cells showed arrest or apoptosis of developing germ cells but does not characterize the immune cells of such tissues (Haimbaugh et al. 2022). In our data, it is unclear whether defects in meiosis or a post-meiotic event is responsible for the absence of these more differentiated cell types in naturally infertile testes. However, the increased presence of immune cells begs the question of whether immune cells act to inhibit fertility or, alternatively, whether they expand in older testes as a response to infertility. If immune cells prevent fertility, one might expect that immunosuppression could rescue fertility in older animals that still contain spermatogonia. Alternatively, immune cells in older testes may regulate spermatogenesis to stave off production of defective sperm. Immune cells may arrive to clear out apoptosing germ cells that were rejected by cell autonomous or hormone-mediated quality control methods (Hikim et al. 2003). Tissue-specific perturbations to the testis immune cell population will be necessary to distinguish between these hypotheses and determine if the increased presence of immune cells in older fish is a cause or consequence of infertility.

Infertile zebrafish testes still contain early germ cells including spermatogonia. It would seem the stem cell pool is not depleted as male zebrafish age, but presence of these cells does not necessarily convey fertility. The spermatogonia population within the testis can be traced back to a small population of embryonic germ cells (Raz 2003). While the spermatogonia population that descends from these embryonic germ cells is maintained in the fertile testis for almost two years, perhaps the clonal diversity declines with age concomitant with the accumulation of detrimental mutations. Lineage tracing of the formation and maintenance of the spermatogonial population may uncover the mechanisms that direct clonality within the spermatogonial population and whether a decrease in clonal diversity accompanies infertility. We anticipate that transcriptomic resources, like those presented here, will facilitate continued explorations of the mechanisms that regulate the development, maintenance, and break down of the zebrafish testis and serve as an evolutionary comparison point for studies of fertility in other vertebrates (Green et al. 2018, Guo et al. 2018, Jung et al. 2019, Guo et al. 2020, Lau et al. 2020, Yu et al. 2021, Nie et al. 2022, Garcia-Alonso et al. 2022, Liu et al. 2022, Huang et al. 2023).

## Methods

### Zebrafish husbandry

All zebrafish used in this study were housed at the University of Utah CBRZ facility. This research was conducted under the approval of the Office of Institutional Animal Care and Use Committee (IACUC protocol 21-02017) of the University of Utah’s animal care and use program.

### Sample preparation and single-cell RNA sequencing

After euthanasia, testes were dissected and immediately placed in a dissociation solution of 440 μL 1X Hank’s Balanced Salt Solution (HBSS), 50 μL 10% pluronic F-68, and 10 μL 0.26U/mL liberase TM. Tissue was incubated at 37°C for approximately 45 minutes while rotating at 750 rpm in an Eppendorf Thermomixer. The dissociation reaction was stopped by adding 1% bovine serum albumin (BSA) in 500 μL cold HBSS. The cell suspensions were filtered through a 40 μm cell strainer. Cells were pelleted in a 4°C centrifuge at 200g for 5 minutes, washed with 0.5% BSA in 500 μL cold HBSS, pelleted again and resuspended in 0.5% BSA in 500 μL cold HBSS. Single-cell RNA sequencing libraries were prepared by the University of Utah High-Throughput Genomics Shared Resource using the 10X Genomics Next GEM Single Cell 3’ Gene Expression Library prep v3.1 with UDI. Libraries were sequenced with the NovaSeq Reagent Kit v1.5 150x150 bp Sequencing.

### Data processing

Raw sequencing reads were processed by using 10X Genomics Cell Ranger software with the v4.3.2 transcriptome reference (Lawson et al. 2020). Further processing and cell clustering was conducted using Seurat v3 (Stuart et al. 2019). Log normalized gene expression was used for clustering of all atlases. To summarize, the barcode-feature matrices from Cell Ranger were converted to Seurat objects. Quality filtering steps included removing cells with >5% mitochondrial genes. Since rates of transcription in the testes are higher than any other organ, we used an interquartile range calculation to determine a reasonable range for the number of genes per cell. We kept cells with >200 genes but fewer than *n* where *n* represents the largest value of the third quartile of each dataset. We used the integration function in Seurat v3 to integrate data sets which considered anchors determined by canonical correlation analysis. Analysis scripts, data, and a web-based app for exploration (Ouyang et al. 2021), are available at https://github.com/asposato/zebrafish_testis_fertility.

### Trajectory analysis with URD

The fertile testis atlas Seurat object was converted to an URD object for pseudotime analysis (Farrell et al. 2018). We used the recommended knn of 199 (square root of total cell number). Each cell within the germ cell atlas was scored according to the state of differentiation. First, we assigned two clusters with the highest *ddx4* expression as the root cells (spermatogonia) and ran URD. Next, we assigned the cluster with the highest *tssk6* expression (spermatozoa) as the root cells and ran URD. We took the mean of the forward pseudotime and the inverse of the reverse pseudotime score to generate a spermatogenesis differentiation score for every germ cell.

### Histology, RNAscope, and Imaging

Tissue for Hematoxylin and eosin (H&E) staining was processed using standard protocols (Siegfried and Steinfeld et al. 2021). Briefly, freshly dissected tissue was fixed in Davidsons’ Fixative overnight at 4ºC. The tissue was then washed with 70% ethanol before processed for paraffin embedding. H&E staining was carried out on 5 μm thick tissue sections. Tissues used for RNAScope were fixed in 4% paraformaldehyde (PFA) overnight at 4ºC, sucrose treated, embedded in optimal cutting temperature compound (OCT), and cryosectioned into 7 μm thick sections. RNAscope® Multiplex Fluorescent Detection Kit v2 (Ref: 323110) and RNAScope probes for *mpeg1*.*1* (macrophages, Ref: 536171-C2) and *acta2* (vasculature, Ref: 508581-C3) were used as markers to identify cell types. Fluorescent images shown are maximum intensity projections that were tiled and stitched using ZEN Black software. Images of dissected testis tissue were taken with a Leica S9E stereo microscope. Histological images were captured with a Zeiss Axio Scan.Z1 Slide Scanner at 40X magnification. Fluorescent images were captured with a Zeiss 880 AiryScan confocal microscope.

## Acknowledgements

We thank all members of the Gagnon lab for helpful discussions and comments. We thank Kellee Siegfried for help with the H&E staining protocol. We thank CZAR and CBRZ staff, especially Nathan Baker, for excellent zebrafish care. We thank the Cell Imaging, Genomics, and Histology core facilities. This project was supported by National Institutes of Health grants R35GM142950 (JAG), T32HD007491 (ALS), 1ZIAHD008997 (JAF), and T32GM141848 (JMW), by a University of Utah (U of U) College of Science Graduate Innovation Fellowship (ALS), by the Las Vegas Alumni Club Scholarship of the University of Utah Alumni Association (ALS), by the U of U Undergraduate Research Opportunity Program (DRL), by the U of U Summer Program for Undergraduate Research (HLH), and by startup funds from the Henry Eyring Center for Cell & Genome Science (JAG). This research was conducted on the traditional and ancestral homeland of the Shoshone, Paiute, Goshute, and Ute Tribes. We affirm and support the University of Utah’s partnership with Native Nations and Urban Indian communities.

## Figures

**Supplemental Figure 1.**
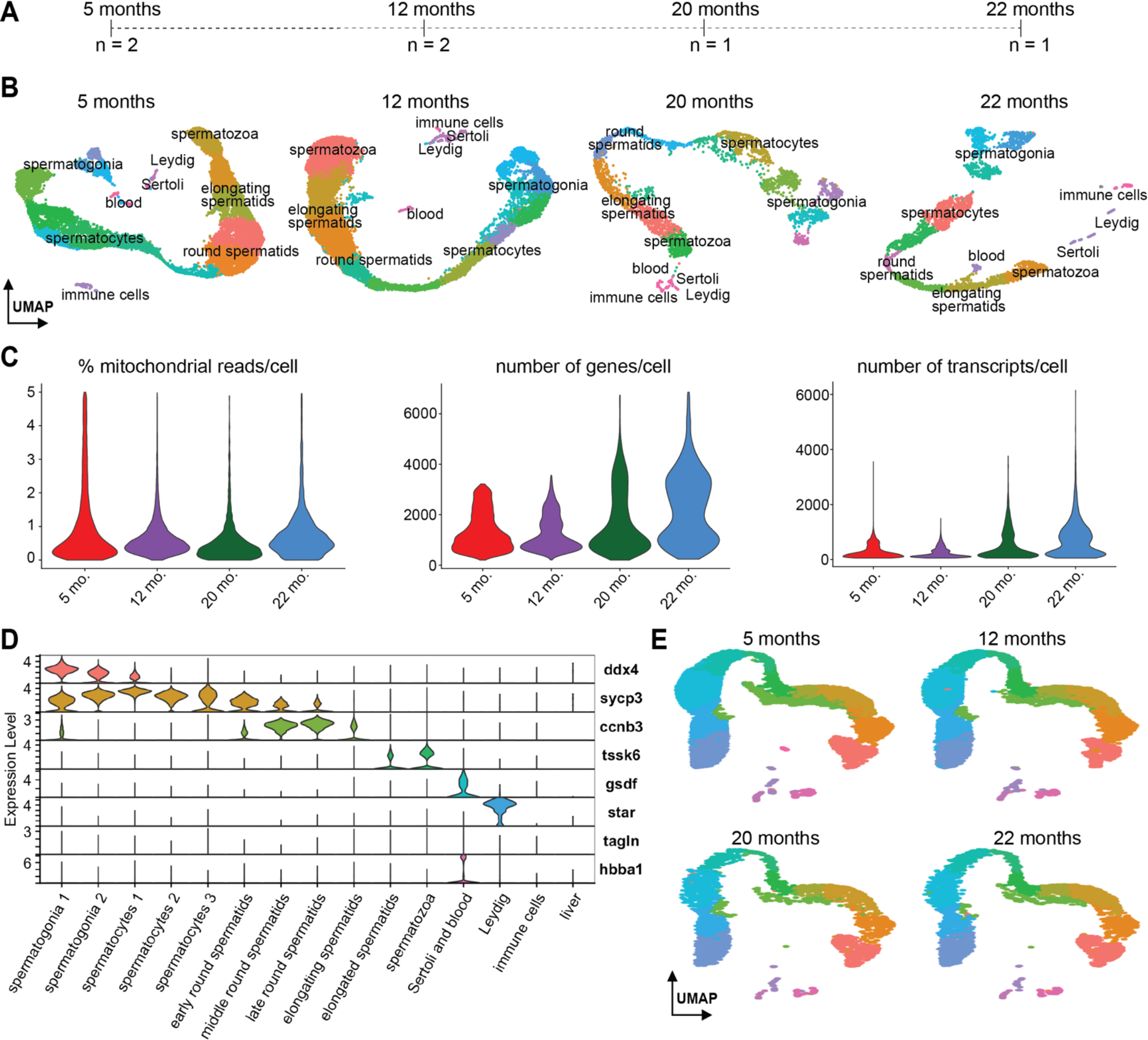
Fertile atlas. **(A)** Ages and sample sizes of testes comprising fertile atlas. **(B)** Individual UMAPs of each age comprising the fertile atlas. **(C)** Violin plots show the percent mitochondrial reads, number of genes, and number of transcripts per cell. **(D)** Violin plots of the marker genes used in Figure 1D-E. **(E)** Fertile atlas split by sample age.

**Supplemental Figure 2.**
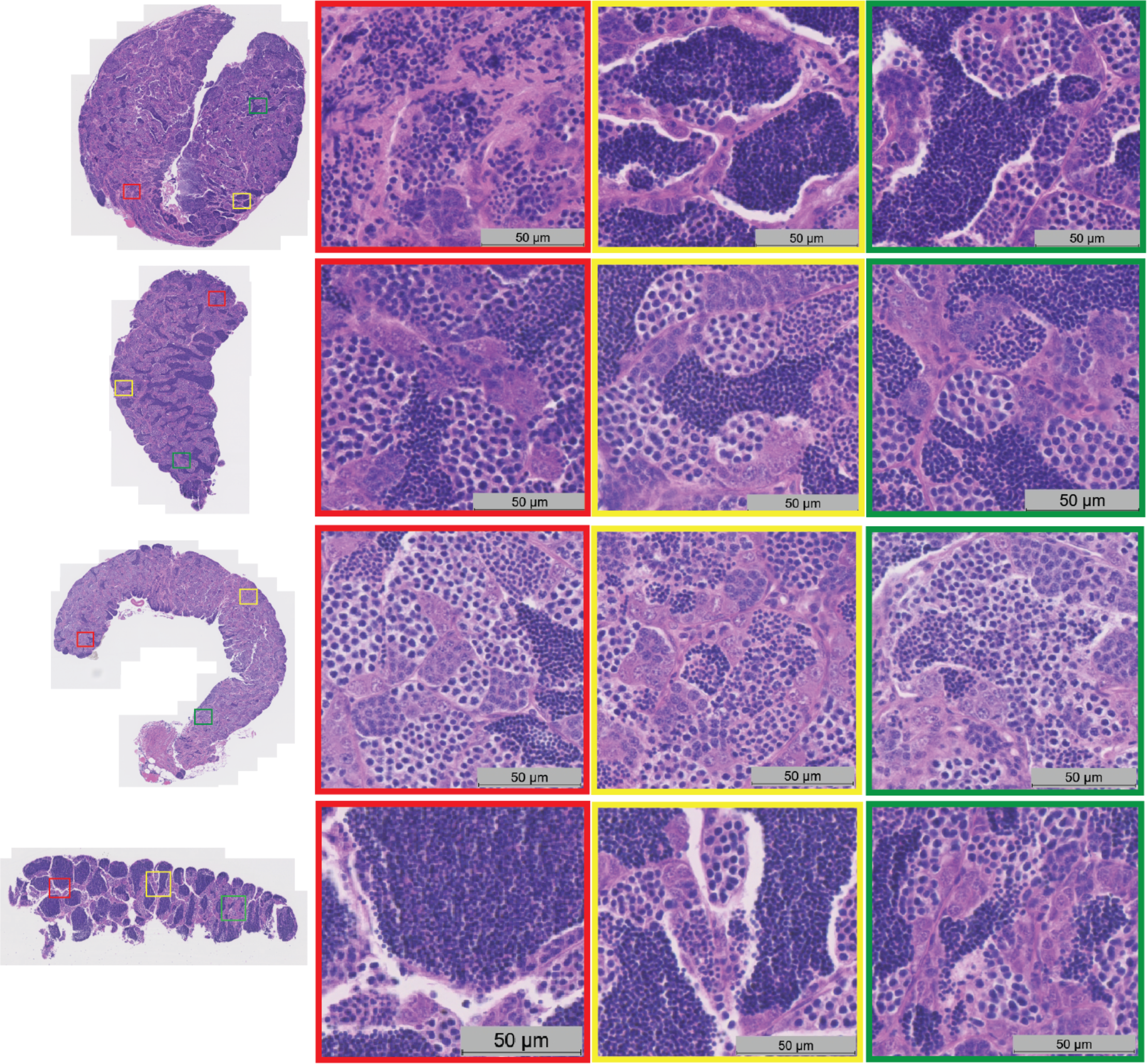
Histology of fertile zebrafish testes.

**Supplemental Figure 3.**
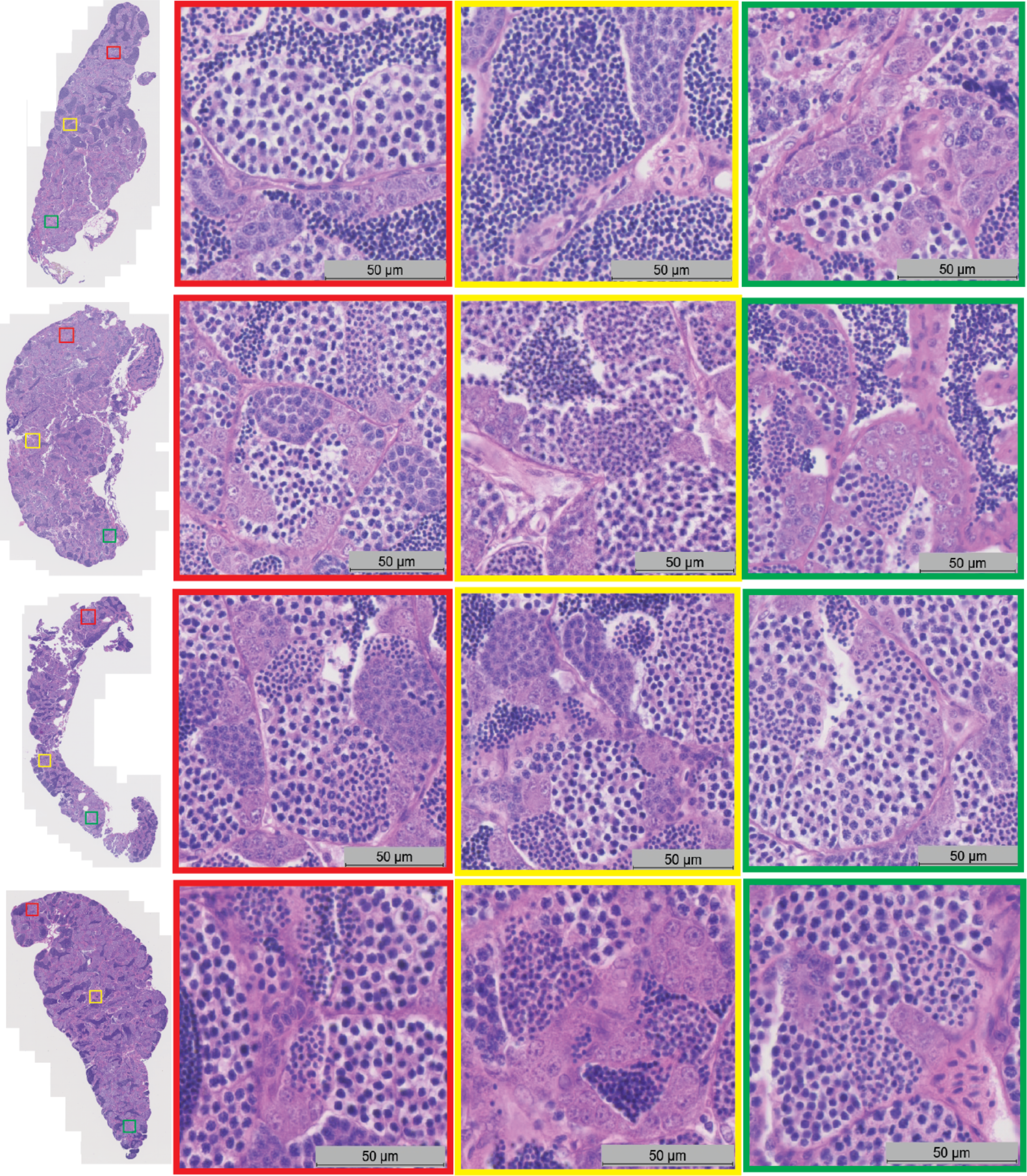
Histology of infertile zebrafish testes.

**Supplemental Figure 4.**
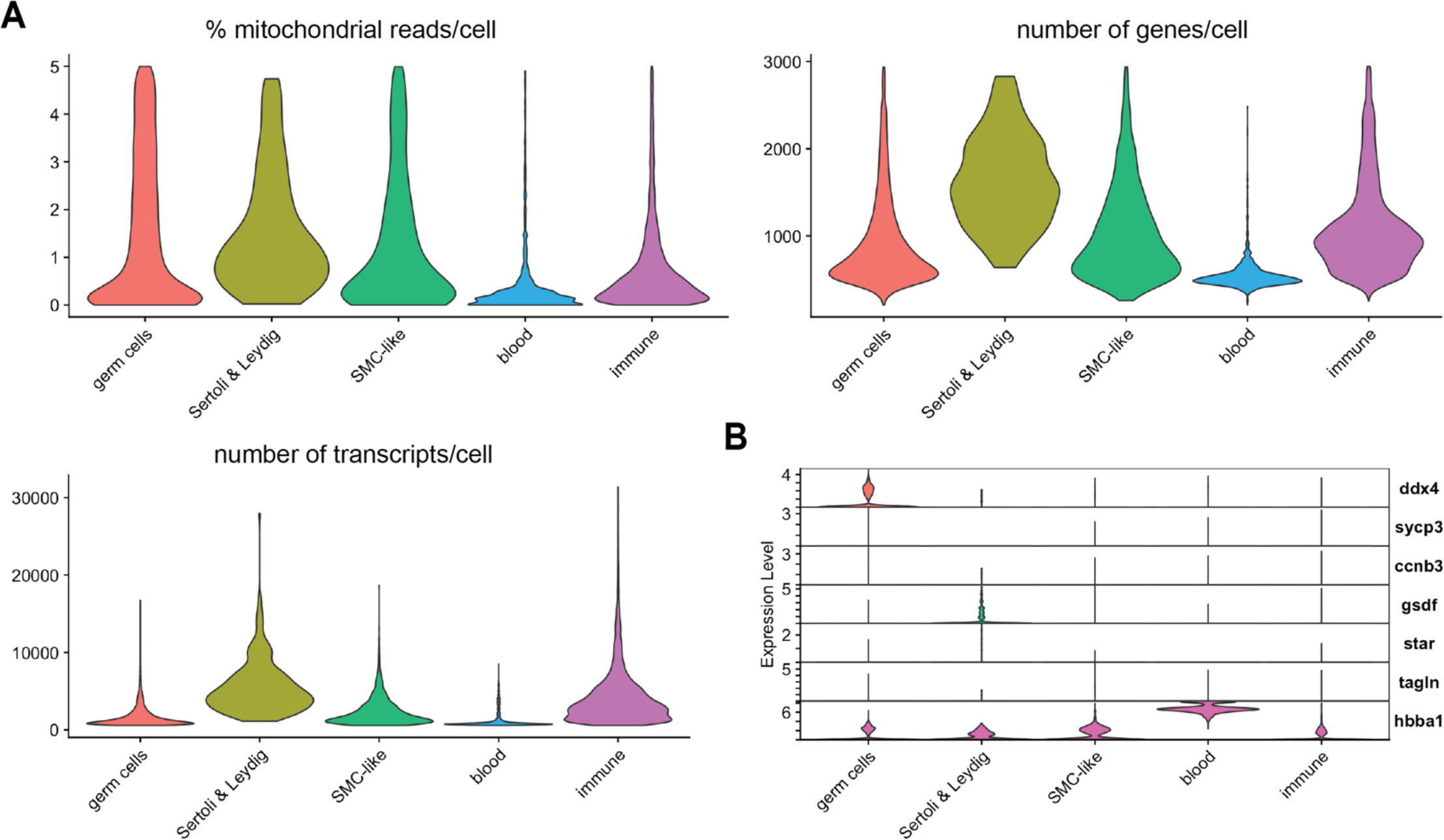
Infertile atlas. **(A)** Violin plots show the percent mitochondrial reads, number of genes, and number of transcripts per cell. **(B)** Violin plots of the marker genes used in Figure 2D-E.

**Supplemental Figure 5.**
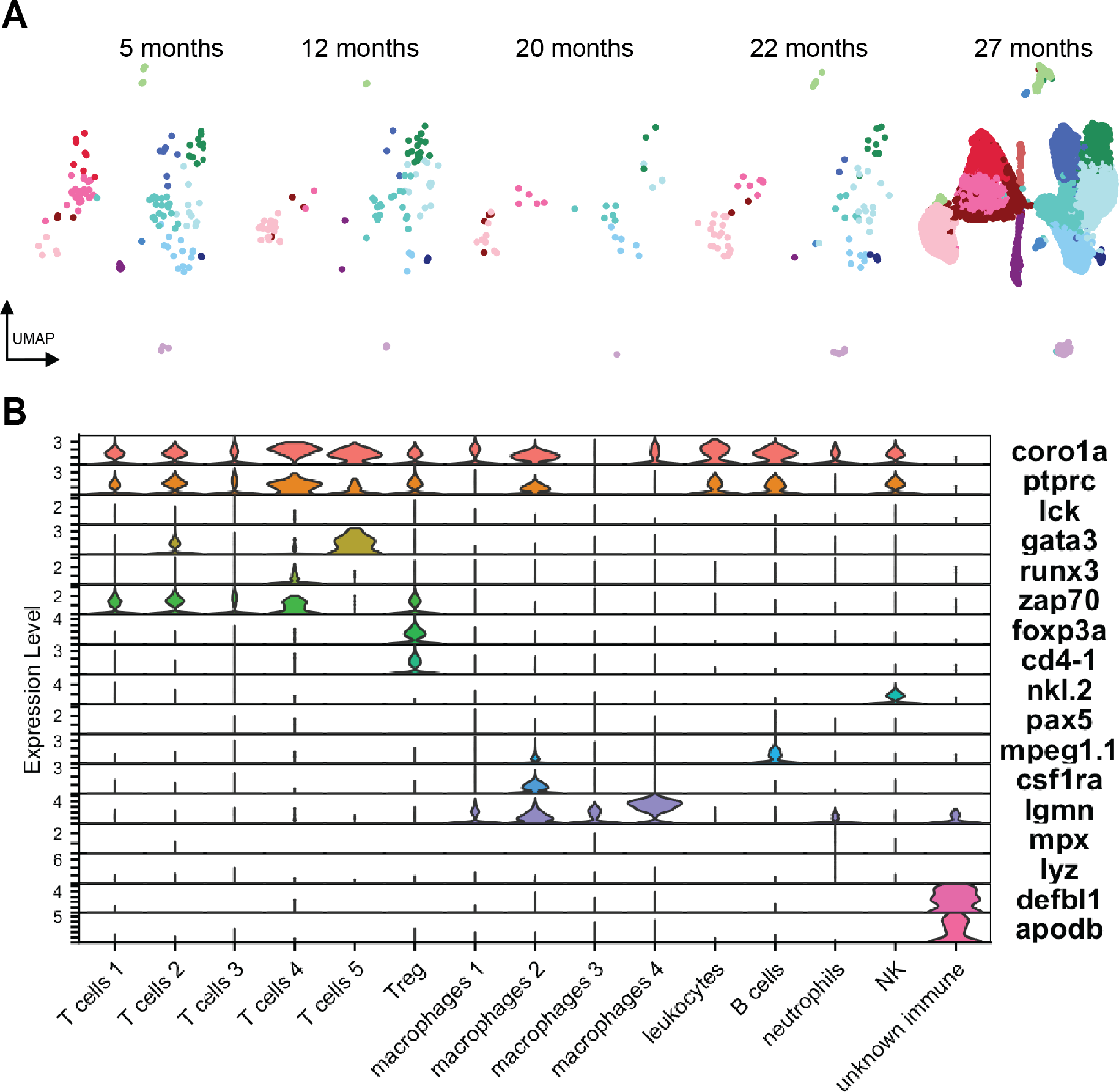
Immune atlas. **(A)** Immune atlas split by sample age. **(B)** Violin plots show marker genes used to identify each cell type.

**Supplemental Figure 6.**
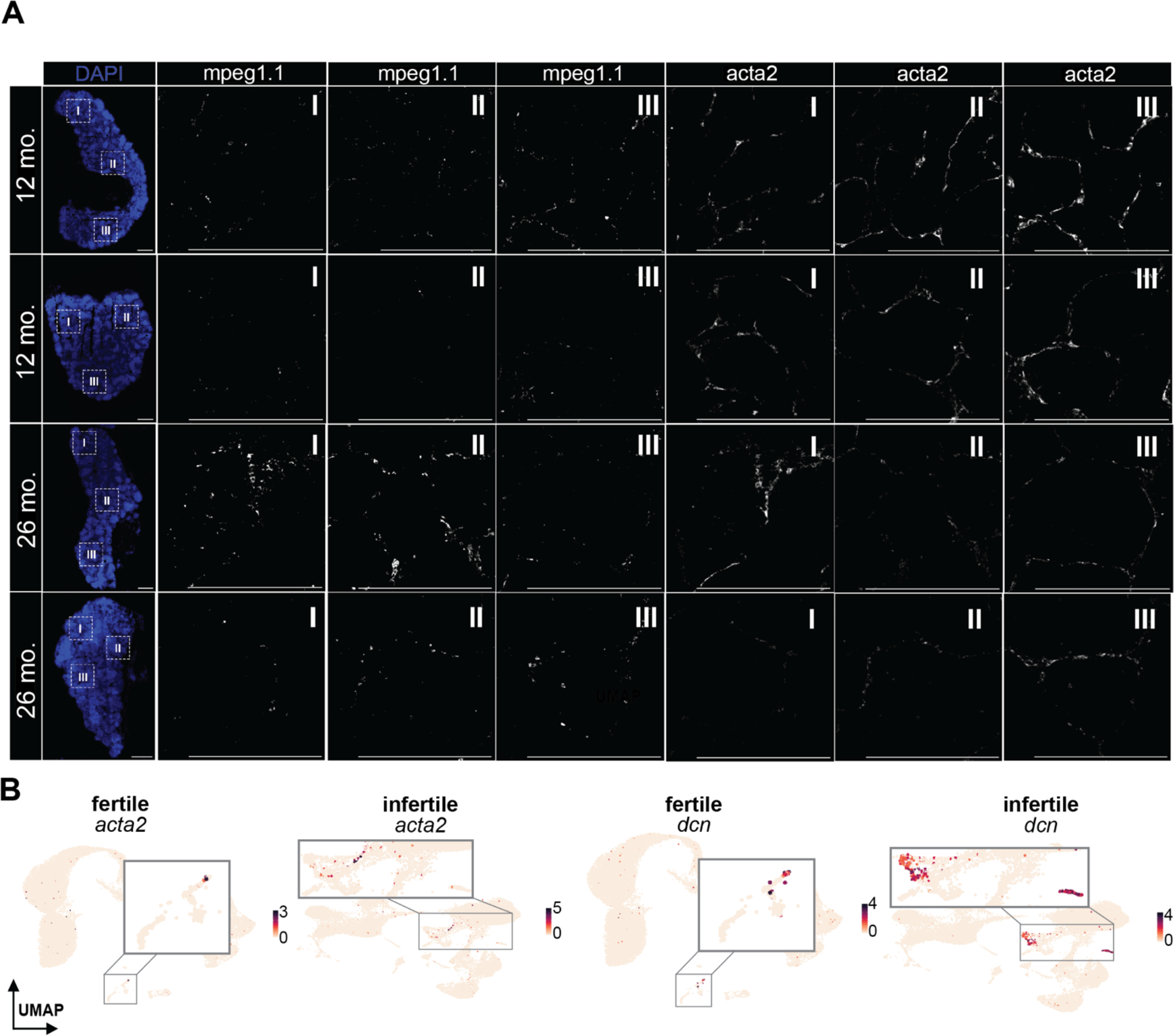
RNA in situ hybridization of immune and smooth muscle cells in fertile and infertile testes. **(A)** RNAscope of fertile and infertile zebrafish testes. **(B)** Expression of markers *acta2* and *dcn* in fertile and infertile atlases. *Acta2* weakly labels smooth muscle cells. *Dcn*, a marker of stressed vascular endothelial cells and fibroblasts, is expressed more abundantly in infertile testes.

**Supplemental Figure 7.**
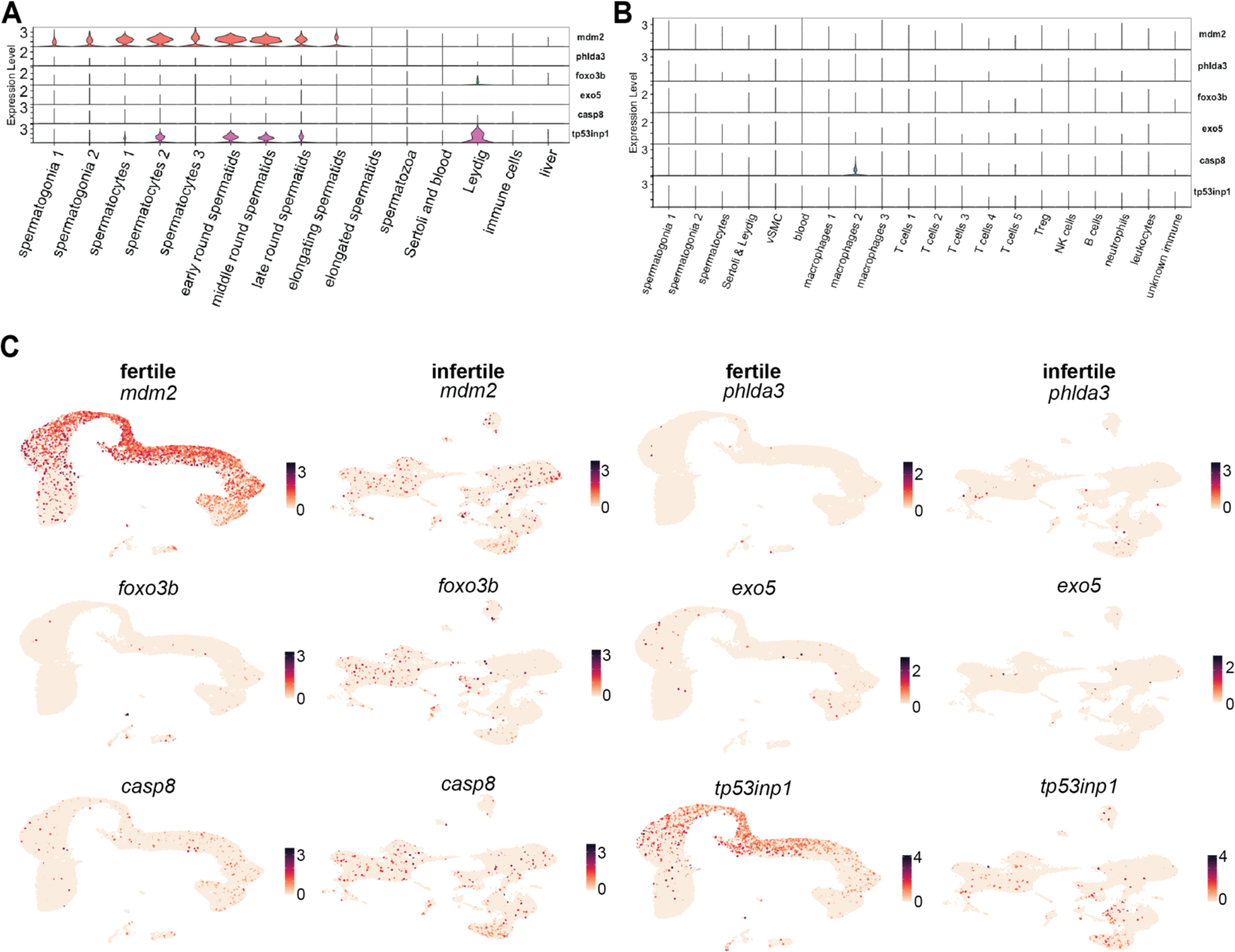
Differences in apoptosis between fertile and infertile testes. (A) Violin plots of apoptotic markers in fertile cell types. (B) Violin plots of apoptotic markers in infertile cell types. (C) Expression patterns of apoptotic markers from A and B in both fertile and infertile testes. All markers positively regulate apoptosis except *mdm2* which is an inhibitor.

